# The history of measles: from a 1912 genome to an antique origin

**DOI:** 10.1101/2019.12.29.889667

**Authors:** Ariane Düx, Sebastian Lequime, Livia Victoria Patrono, Bram Vrancken, Sengül Boral, Jan F. Gogarten, Antonia Hilbig, David Horst, Kevin Merkel, Baptiste Prepoint, Sabine Santibanez, Jasmin Schlotterbeck, Marc A. Suchard, Markus Ulrich, Navena Widulin, Annette Mankertz, Fabian H. Leendertz, Kyle Harper, Thomas Schnalke, Philippe Lemey, Sébastien Calvignac-Spencer

**Affiliations:** Epidemiology of Highly Pathogenic Microorganisms, Robert Koch Institute, Berlin, Germany; Viral Evolution, Robert Koch Institute, Berlin, Germany; KU Leuven Department of Microbiology, Immunology and Transplantation, Rega Institute, Laboratory of Clinical and Evolutionary Virology, Leuven, Belgium; Institute for Pathology, Charité, Berlin, Germany; National Reference Centre Measles, Mumps, Rubella, Robert Koch Institute, Berlin, Germany; Department of Biostatistics, Fielding School of Public Health and Departments of Biomathematics and of Human Genetics, David Geffen School of Medicine, University of California, Los Angeles, CA, USA; Berlin Museum of Medical History at the Charité, Berlin, Germany; Department of Classics and Letters, The University of Oklahoma, Norman, OK, USA

## Abstract

Many infectious diseases are thought to have emerged in humans after the Neolithic revolution. While it is broadly accepted that this also applies to measles, the exact date of emergence for this disease is controversial. Here, we sequenced the genome of a 1912 measles virus and used selection-aware molecular clock modeling to determine the divergence date of measles virus and rinderpest virus. This divergence date represents the earliest possible date for the establishment of measles in human populations. Our analyses show that the measles virus potentially arose as early as the 4^th^ century BCE, rekindling the recently challenged hypothesis of an antique origin of this disease.

**One Sentence Summary:** Measles virus diverged from rinderpest virus in the 4^th^ century BCE, which is compatible with an emergence of measles during Antiquity.

---

Measles is a highly contagious viral disease that presents with rash, fever and respiratory symptoms. Before a live-attenuated vaccine was developed in the 1960s, the disease affected the vast majority of children (*1*, *2*). Global vaccination campaigns resulted in a marked reduction of measles transmission and fatal cases and WHO has proclaimed an elimination goal. However, the disease still caused an estimated 110,000 deaths in 2017 (*3*) and incidence has recently been on the rise (*4*). Measles is caused by *Measles morbillivirus* (MeV), a negative sense single-stranded RNA virus from the family *Paramyxoviridae* (Order: *Mononegavirales*). MeV is an exclusively human pathogen whose closest relative was the now eradicated *Rinderpest morbillivirus* (RPV), a devastating cattle pathogen (*5*). It is generally accepted that measles emergence resulted from a spill-over from cattle to humans, although the directionality of this cross-species transmission event has never been formally established (supplementary text S1, *6*).

It is unclear when measles first became endemic in human populations, but assuming an origin in cattle, the earliest possible date of MeV emergence is defined by the MeV-RPV divergence time. Several studies have provided estimates for this date using molecular clock analyses (*7*–*10*), with the most reliable (and oldest) estimate falling at the end of the 9^th^ century CE (mean: 899 CE; 95% highest posterior density (HPD) interval: 597 – 1144 CE) (*8*). Here, we reassess the MeV-RPV divergence time using advanced, selection-aware Bayesian molecular clock modelling (*11*) on a dataset of heterochronous MeV genomes including the oldest human RNA virus genome sequenced to date. We show that a considerably earlier emergence during Antiquity can no longer be excluded.

Our re-examination was prompted by the broadly accepted view that molecular dating based on tip date calibration, i.e. the method used in previous efforts to estimate the timing of MeV-RPV divergence, underestimates deep divergence times (*8*). Rapid short-term substitution rates captured by tip calibration can often not be applied over long evolutionary timescales, because of the effects of long-term purifying selection and substitution saturation. This causes a discrepancy between short- and long-term substitution rates, which is referred to as the time-dependent rate phenomenon (TDRP; *12*, *13*). Since measurement timescales matter, a first step to arrive at accurate estimates is to maximize the time depth of tip calibration, for example through the use of ancient viral sequences (*14*, *15*).

RNA tends to be much less stable in the environment than DNA, making the recovery of MeV genetic material from archeological remains unlikely (*16*, *17*). Pathology collections represent a more realistic source of MeV sequences that predate the oldest MeV genome – the genome of the Edmonston strain that was isolated in 1954 and attenuated to become the first measles vaccine. We examined a collection of lung specimens gathered by Rudolf Virchow and his successors between the 1870s and 1930s and preserved by the Berlin Museum of Medical History at the Charité (Berlin, Germany), and identified a 1912 case diagnosed with fatal measles-related bronchopneumonia (Fig. 1, fig. S1, supplementary texts S2 and S3). To retrieve MeV genetic material from this specimen, we first heat treated 200mg of the formalin-fixed lung tissue to reverse macromolecule cross-links induced by formalin and subsequently performed nucleic acid extraction (*18*). Following DNase treatment and ribosomal RNA depletion, we built high-throughput sequencing libraries and shotgun sequenced them on Illumina® platforms. We generated 27,328,219 reads, of which 0.46% were mapped to MeV genome, representing 10,960 unique MeV reads. These allowed us to reconstruct an almost complete 1912 MeV genome: 15,257 of the 15,894 nucleotides in the MeV strain Edmonston (AF266288) were covered by at least 3 unique reads (11,988 nucleotides by at least 20 reads; mean coverage 54x).

**Fig. 1.**
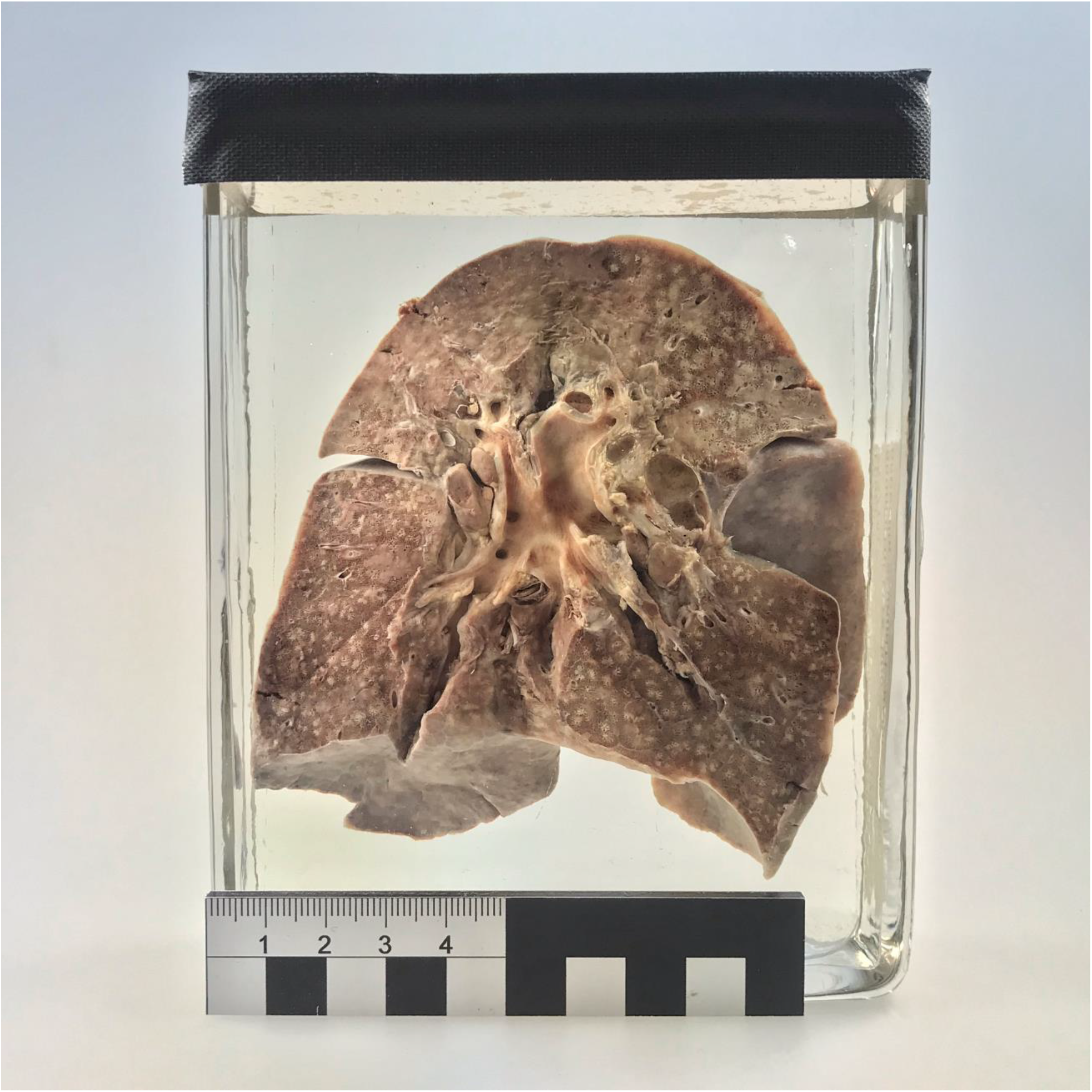
Formalin-fixed lung specimen collected in 1912 in Berlin from a 2-year old girl diagnosed with measles-related bronchopneumonia (museum object ID: BMM 655/1912).

In addition to the 1912 genome and the 1954 Edmonston genome, only 2 genomes have been determined from MeV isolated prior to 1990 (Mvi/Lyon.FRA/77: HM562899; T11wild: AB481087). We therefore searched the strain collection of the German National Reference Laboratory (Robert Koch Institute, Berlin, Germany) for pre-1990 isolates. We found two strains from the pre-vaccine era isolated in 1960 by the National Reference Laboratory of former Czechoslovakia in Prague (MVi/Prague.CZE/60/1 and MVi/Prague.CZE/60/2; *19*). We performed serial passages of these strains and determined their genome sequences at a mean coverage of 109x and 70x, respectively. The two genomes were nearly identical, differing at only four sites.

We performed Bayesian and maximum likelihood (ML) phylogenetic analyses to investigate the phylogenetic placement of the 1912 and 1960 genomes with respect to 127 available MeV genomes. Tip-dated Bayesian phylogenetic trees placed the 1912 genome basal to all modern genomes while the two genomes from 1960 clustered together with the Edmonston strain (genotype A; fig. S2). The basal placement of the 1912 genome was further confirmed by exploring the rooting of non-clock ML trees based on the relationship between sampling time and root-to-tip-divergence (both including and excluding this genome; supplementary text S4 and fig. S3 A and B). The relatedness of the 1912 and 1960 genomes to now extinct MeV lineages is in line with a marked reduction of MeV genetic diversity during the 20^th^ century as a product of massive vaccination efforts.

Having extended the time depth of MeV tip calibration, we subsequently focused our attention on estimating the timing of MeV-RPV divergence. We assembled a dataset of 51 genomes comprising MeV (including one of the 1960 genomes and the 1912 genome), RPV and Peste des petits ruminants virus (PPRV, the closest relative to MeV-RPV) sequences, ensuring they represented the known genetic diversity of these viruses. This dataset was used to infer a time-scaled evolutionary history for these three species in a Bayesian phylogenetic framework. We constructed a series of evolutionary models with increasing degree of complexity to accommodate various sources of rate heterogeneity. Models ranged from a standard codon substitution model with a strict molecular clock assumption to a codon substitution model with time-varying selection pressure combined with a clade-specific rate for PPRV and additional branch-specific random effects on the substitution rate. Adequately accommodating different sources of rate heterogeneity is known to provide a better correction for multiple hits in genetic distance estimation and the potential of codon substitution modelling in recovering deep viral divergence has specifically been demonstrated (*8*). This was reflected in the increasingly older estimates of MeV-RPV and PPRV-MeV-RPV divergence times and wider 95% highest posterior density intervals in increasingly complex models (Fig. 2 and table S2). Parameter estimates of the substitution and clock models also provided evidence for a significant contribution of these different sources of rate heterogeneity to model fit improvement (table S2, supplemetary text S4). We found a significantly negative coefficient for the time-dependent nonsynonymous/synonymous substitution rate ratio (ω) (*11*), indicating strong long-term purifying selection, a significantly positive coefficient for the fixed effect on the PPRV rate, indicating a faster evolutionary rate in this clade (as suggested by temporal signal analyses, fig. S4), and significant additional unexplained variation as modelled by the random effects. Our most complex model therefore provided the best description of the evolutionary process and pushed back the divergence date of MeV and RPV to Antiquity, with a mean estimate at 345 BCE [95% highest posterior density (HPD) interval: 1066 BCE - 319 CE] (Fig. 3A). These estimates were robust to including or estimating the age of the 1912 genome in the analyses; estimating its sampling date resulted in a posterior distribution in agreement with the known date (1923 CE [95% HPD interval: 1883 - 1962 CE]; supplementary text S5 and table S2).

**Fig. 2.**
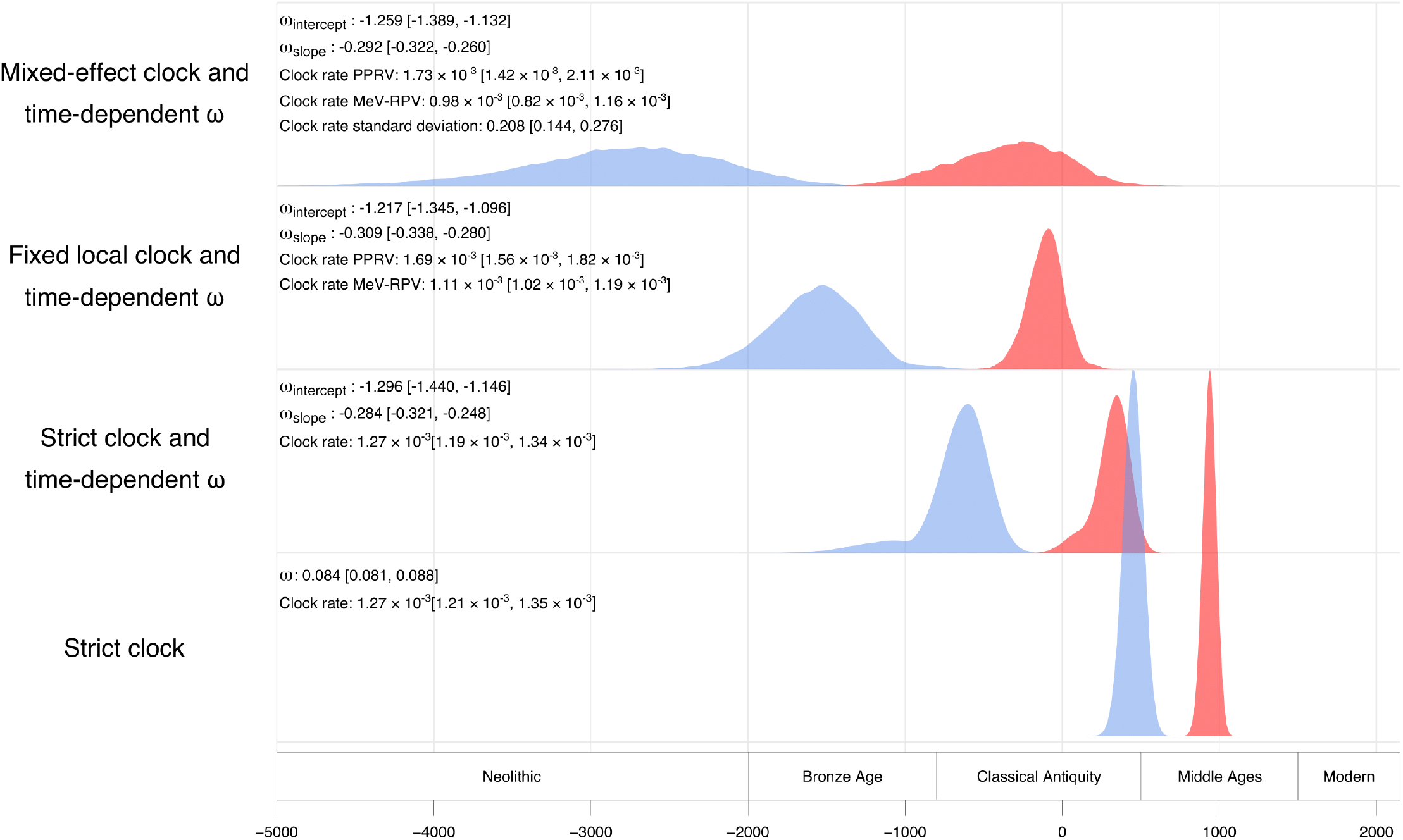
Influence of model complexity on the divergence time estimates between MeV and RPV (red), and between MeV/RPV and PPRV (blue). Estimated parameter values (posterior mean and 95% highest posterior density interval) for each model are provided on the left. Historical divisions are based on classical “world history” periods.

**Fig. 3.**
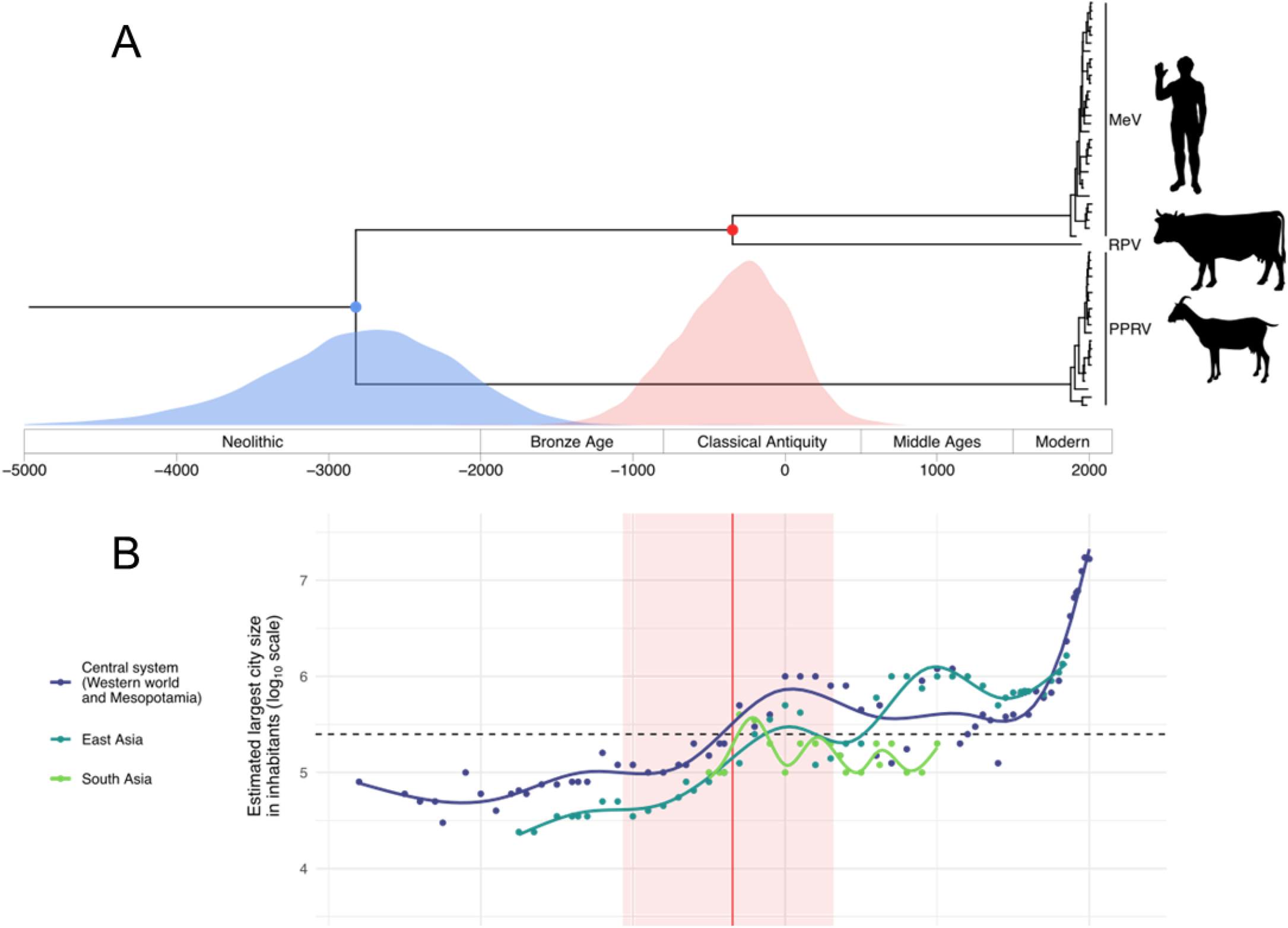
Estimation of the divergence time between MeV and RPV, and between MeV/RPV and PPRV (A) and estimated size (log10 scale) of the largest city in the Central system (dark blue), East Asia (petrol) and South Asia (green) over time (B). (A) Maximum clade credibility (MCC) tree summarized from a Bayesian time-measured inference using tip-dating and accounting for long-term purifying selection. The red and blue points represent the divergence events of MeV and RPV and MeV/RPV and PPRV, respectively; the according divergence date estimates are reported below as marginal posterior distributions. Historical divisions are based on classical “world history” periods. (B) The red vertical line represents the divergence time estimate between MeV and RPV and the red area its 95% highest posterior density interval. The dashed horizontal line represents the classical threshold for MeV maintenance in a population (i.e. 250,000 individuals). Dots show data points according to Morris (*29*) and Inoue et al. (*26*). Each line represents the fit of a generalized additive model with a cubic spline smoothing function.

While the divergence of MeV and RPV *predates* the emergence of measles in humans, it does not necessarily coincide with it. This new estimate should therefore be considered a lower bound for measles emergence, which is now compatible with the emergence of this disease during Antiquity. This raises the question of whether an earlier timing of measles emergence agrees with other sources of information.

The earliest clear clinical description of measles is often attributed to the Persian physician Rhazes, writing in the 10^th^ century CE (*20*). But Rhazes was extremely familiar with all available medical literature at his time, and made use of earlier sources. Indian medical texts probably describe measles several centuries prior to Rhazes (*21*). While clear descriptions of measles are missing in the Hippocratic corpus and the Greek medical tradition (at least through the prolific second-century writer Galen), such absence alone cannot be decisive. Retrospective diagnosis from pre-modern medical texts is notoriously fraught, especially for diseases like measles whose symptoms were easily confused with a variety of other conditions. Measles differential diagnosis remained a challenge well into more recent times (*22*). Therefore, any number of the large-scale “pestilences” described in first-millennium sources from Europe or China could reflect MeV outbreaks.

An antique origin of measles seems all the more plausible in the light of demographic changes that are compatible with our understanding of MeV (contemporary) epidemiology. Indeed, Antiquity witnessed an important upsurge in population sizes in Eurasia and South and East Asia. Populations large enough to support continuous MeV transmission, i.e. larger than the MeV critical community size (CCS) of 250,000-500,000 individuals (*23*–*25*), have existed in Eurasia since the first millennium BCE. Although population sizes derived from ancient documents or archaeological proxies are only estimates (supplementary text S6), there is broad agreement that a number of settlements in North Africa, India, China, Europe, and the Near East began to surpass the CCS for MeV by around 300 BCE, presumably for the first time in human history (Fig. 3B; *26*).

Based on these considerations, our substantially older divergence estimate provides grounds for sketching a new model of MeV’s evolutionary history. Under this scenario, a bovine virus, the common ancestor to modern strains of RPV and MeV, circulated in large populations of cattle (and possibly wild ungulates) since its divergence from PPRV around the 3rd millennium BCE (mean estimate at 2821 BCE [95% HPD interval: 4177 - 1665 BCE]; Fig. 3A). As a fast-evolving RNA virus, it may have produced variants that were able to cross the species barrier on several occasions, but small human populations could only serve as dead-end hosts. Then, almost as soon as contiguous settlements reached sufficient sizes to maintain the virus’ continuous transmission (Fig. 3B), it emerged as a human pathogen, the progenitor of modern-day MeV. Concurrent human-bovine epidemics that are well documented in e.g. Roman sources from the 5^th^ century BCE on (*27*) may mark the early stages of MeV emergence when the ancestor of MeV was pathogenic to both cattle and humans. During the following centuries, introduction of MeV into naive populations and/or flare-ups of the disease might have caused some ancient epidemics whose etiology remains uncertain.

While our findings shed new light on the origin of measles, formally proving that the virus emerged during Antiquity would require archeological genomic evidence. Recently, a 1000-year-old plant RNA virus genome (Zea mays chrysovirus 1) was recovered from maize remains (*28*) and genetic material of parvovirus B19 was detected in early Neolithic skeletal remains, despite the relatively unstable nature of its single-stranded DNA genome (*15*). Such advances highlight that it may not be completely impossible for antique remains to still contain MeV RNA, especially if preserved under favorable circumstances, including natural mummification or preservation in cold environments (*16*, *17*). While awaiting such direct evidence, we believe that the proposed model of MeV evolution constitutes a compelling working hypothesis.

## Acknowledgments

The Titan V GPU used for this research was donated by the NVIDIA Corporation.

## Author contributions

Conceptualization: PL, SCS; data curation: AD, SL, BV, MAS, PL, SCS; formal analysis: AD, SL, BV, JFG, MU, PL, SCS; funding acquisition: FHL, SCS; investigation: AD, LVP, SB, AH, DH, KM, BP and JS; methodology: SL, BV, MAS, PL; project administration: FHL, PL, SCS; resources: SS, NW, AM, FHL, TS, PL, SCS; software: SL, BV, MAS, PL; supervision: PL, SCS; validation: AD, MU, JFG, SCS; visualization: SL; writing – original draft: AD, SL, KH, PL, SCS; writing – review and editing: all authors.

## Data and materials availability

MeV sequences from 1912 and 1960 are deposited in GenBank under XXX. Alignments, trees, and BEAST xml files are available at XXX.

