## Supplemental material for "The history of measles: from a 1912 genome to an antique origin"

**This PDF file includes:**

Materials and Methods

Supplementary Text S1 to S6

Figures S1 to S5

Tables S1 to S2

References *30* to *69*

### Materials and Methods

#### Ethics

Ethics approval was obtained from the ethics committee of the Charité (Berlin, Germany) under the reference number EA4/212/19.

#### Precautions for ancient DNA

To avoid contamination with modern DNA, all laboratory analyses preceding amplification for dual indexing were performed in the clean room of our ancient DNA laboratory. Here, specific precautions are applied, including restricted entry to trained personnel wearing full body suits, masks, overshoes and double gloves, and daily UV light decontamination of the whole room. Prior to this study, no measles positive samples were handled in these facilities.

#### Sample material

We obtained a sample of a formalin-fixed lung specimen (museum object ID: BMM 655/1912) from the collections of the Berlin Museum of Medical History at the Charité (Berlin, Germany). The specimen is a lung that was collected during an autopsy of a 1912 measles patient from Berlin. According to the case file, the lung belonged to a 2-year-old child that died on June 3$^{rd}$, 1912, of measles-related bronchopneumonia. Since the exact composition and concentration of the formalin fixative was unknown, we stored the collected lung sample in PBS to avoid further damage to nucleic acids by adding fresh formalin.

In addition to the 1912 measles case, we obtained two *Measles morbillivirus* (MeV) strains (MVi/Prague.CZE/60/1 and MVi/Prague.CZE/60/2) which were isolated by the National Reference Laboratory of former Czechoslovakia in Prague in 1960. These strains were kept by the National Reference Laboratory of the former German Democratic Republic until they became part of the strain collection of the German National Reference Laboratory (Robert Koch Institute, Berlin, Germany) after the German reunification. The strains were isolated in the intraamniotic cavity of chick embryos and passaged inVero cells for 5 and 6 passages, respectively, and for one passage in human SLAM-expressing Vero cells.

Extraction

To maximize chances of viral RNA recovery, we performed eight separate nucleic acid extractions from different areas of the lung using the DNeasy® Blood & Tissue Kit (Qiagen) with modifications for formalin-fixed samples. For each separate extraction, a pea-sized piece (ca. 25mg) of lung was washed in 1 ml PBS to remove residual fixative. The washed tissue was cut into smaller pieces using sterile scissors and added to bead tubes containing tissue lysis buffer (ATL). To reverse formalin-induced crosslinking, the tissue was heated to 98°C for 15 minutes. To facilitate lysis, the tissue was homogenized by bead beating for three times 20s at 4m/s with a Fast Prep® (MP Biomedicals). We added 20 µl Proteinase K and kept the homogenate at 56°C until the tissue was completely lysed (ca. 1 hour). Subsequent steps were performed according to the manufacturer's protocol and nucleic acids were eluted in 35 µl elution buffer (AE).

Library preparation

To maximize viral RNA content in the final sequencing libraries, we removed DNA and ribosomal RNA from the nucleic acid extracts before conversion to double-stranded cDNA. For DNase treatment we used the TURBO DNA-*free*™ Kit (Ambion). We performed ribosomal RNA depletion and clean-up separately on the first four DNAse-treated extracts, using NEBNext® rRNA Depletion Kit (Human/Mouse/Rat) with RNA Sample Purification Beads (New England Biolabs) according to protocol. To reduce costs, we pooled and concentrated the next four DNAse-treated extracts using RNA Clean & Concentrator-5 Kit (Zymo Research) and eluted in 13 µl nuclease free water as input for one ribosomal RNA depletion reaction. In both cases, following bead clean-up, RNA was eluted in 12 µl nuclease free water. We performed cDNA synthesis, using the SuperScript™ IV First-Strand Synthesis System (Invitrogen) and converted cDNA into dsDNA with the NEBNEXT® mRNA Second Strand Synthesis Module (New England Biolabs). Double-stranded DNA was purified using MagSi-NGS$^{prep}$ Plus Beads (Steinbrenner Laborsysteme) and eluted in 50 µl Tris-HCl (10mM) EDTA (1mM) Tween20 (0.05%) (TET) buffer. We prepared 5 separate libraries with the NEBNext® Ultra™ II DNA Library Prep Kit for Illumina® (New England Biolabs) without prior fragmentation of double-stranded cDNA and without size-selection upon adapter ligation. All clean-up steps during the library preparation were conducted with MagSi-NGS$^{prep}$ Plus Beads (Steinbrenner Laborsysteme). Libraries were dual indexed with NEBNext® Multiplex Oligos for Illumina® (New England Biolabs), quantified using the KAPA Library Quantification Illumina Universal Kit (Roche), amplified with the KAPA HiFi HotStart ReadyMix (Roche) and Illumina adapter-specific primers, and diluted to a concentration of 4nM for sequencing.

Sequencing

Libraries were sequenced on an Illumina® MiSeq platform using the v3 chemistry (2x300-cycle) and on an Illumina® NextSeq platform using v2 chemistry (2x150-cycle) for a total of 48,326,978 unfiltered reads.

Filtering and genome assembly

Sequencing reads were filtered (adapter removal and quality filtering) using Trimmomatic (*30*) with the following settings: LEADING:30 TRAILING:30 SLIDINGWINDOW:4:30

MINLEN:30. For the 1912 sequences, we attempted *de novo* assembly on a subsample of 2,083,813 reads (up to 500,000 reads per library) using metaSPAdes (*31*). However, since the largest MeV contig generated with this approach covered only 1657 nt, we chose to use a two-step reference based mapping approach. We first used BWA-MEM (*32*) to map trimmed reads to the MeV RefSeq genome (NC_001498). We then called the consensus of the two largest contigs and determined the closest MeV genome via BLASTn search against the NCBI non-redundant nucleotide collection (*33*). The best hit for both contigs was the Measles virus strain Edmonston (AF266288). Subsequently, we merged paired reads with ClipAndMerge (*34*) and used BWA-MEM to map all reads (merged, unmerged, unpaired) to the Edmonston genome. We sorted the mapping files and removed duplicates with the tools SortSam and MarkDuplicates from the Picard suite (*35*). We compared MarkDuplicates to Dedup (*36*) to select the more stringent tool for duplicate removal, finding that for all libraries MarkDuplicates removed more reads. We proceeded with these more conservative mapping results. For the final genome assembly, we merged the mapped reads (bam files) of the five libraries using MergeSamFiles from the Picard suite (*35*). Using Geneious 11.1.5 (*37*), we assembled two consensus genomes, 1) with a minimum coverage of 20x and a 95% majority for base calling (modern DNA settings), and 2) with a minimum 3x coverage and 50% majority consensus (ancient DNA settings). The consensus sequences contained 11,988 and 15,257 unambiguous positions, respectively. The 1960s MeV strains were treated in the same way resulting in consensus of 15,870 nts and 15,868 nts (minimum 3x coverage; 50% majority consensus), and 15,780 nts and 15,713 nts (minimum 20x coverage; 95% majority consensus), respectively.

<u>Phylogenetic analyses</u>

<u>Dataset preparation</u>

We collected all available full length measles (MeV), rinderpest (RPV) and "peste des petits ruminants" (PPRV) virus genomes from GenBank, and excluded vaccine strains and subacute sclerosing panencephalitis (SSPE) MeV strains. This resulted in a dataset of 133 MeV (including the 3 new sequences from this study), 1 RPV, and 73 PPRV full-length genomes. Sequences from MeV and PPRV were aligned independently using MAFFT v7 (*38*) and checked for recombination using RDP4 (*39*). Sequences exhibiting a significant result for more than three of the selected recombination detection methods (RDP, GENECONV, Chimaera, MaxChi, Bootscan, SiScan, and 3Seq) were excluded from subsequent analyses. The final dataset consisted of 130 MeV, 65 PPRV and 1 RPV full-length genomes.

#### Preliminary phylogenetic analysis

Phylogenetic trees for the final full-length MeV and PPRV genomes were reconstructed using IQ-TREE version 1.6.10 (*40*) under a general time-reversible (GTR) substitution model with discrete Γ-distributed rate variation among sites. We used TempEst v1.5.1 (*41*) to visually explore the temporal signal in the resulting maximum likelihood trees by plotting root-to-tip divergence as a function of sampling time with the root that optimized their correlation for both MeV and PPRV.

#### Phylogenetic analysis of the 1912 sample

A full phylogenetic tree with the 130 full-length MeV genomes was constructed using BEAST v1.10.4 (*42*) under a GTR + Γ nucleotide substitution model, an uncorrelated relaxed lognormal clock and a constant population size. Multiple independent MCMC chains were run sufficiently long to ensure proper mixing after convergence to the posterior of the model parameters, as measured by sufficiently high effective sample sizes (ESS > 200) using Tracer v1.7 (*43*). Trees and log files were combined after removing burn-in and posterior distribution of trees was summarized as a maximum clade credibility (MCC) trees using TreeAnnotator 1.10.4 (fig. S1; *42*).

To confirm and explore to which extent the basal position of the 1912 sample was driven by its age, we inferred a maximum likelihood (ML) tree for the remaining 130 full-length MeV genomes in IQ-TREE version 1.6.10 (*40*) under a general time-reversible (GTR) and discrete Γ model. Another tree was generated under the same conditions without the 1912 isolate genome. The impact on the temporal signal of the 1912 sample was explored by computing the correlation between sampling time and root-to-tip divergence: while the tree rooting was optimized with the 1912 genome sequence, the correlation was computed without it.

Phylogenetic analyses to date RPV-MeV divergence

We inferred maximum likelihood (ML) trees for the remaining 130 MeV and 65 PPRV full-length genomes in IQ-TREE version 1.6.10 (*40*) under a general time-reversible (GTR) and discrete Γ model. These were used to subsample the dataset while keeping an optimal representation of the genetic diversity of each virus using Phylogenetic Diversity Analyzer v0.5 (*44*). This step was necessary to ensure manageable run-time during subsequent analysis with complex codon models in a Bayesian framework. The final set (table S1) consisted of 30 MeV genomes, 20 PPRV genomes and 1 RPV genome, including two out of the three new sequences generated in this study.

All 51 full-length genomes were aligned using MAFFT and trimmed to keep only conserved coding regions among the three viral species. The alignment was manually checked at the codon level to ensure all sequences retained a proper translation frame; one sequence (accession number GQ376026) had an improper coding frame that could be corrected by inserting an N at position

7,025, resulting in a correct amino-acid sequence, similar to other isolates of the same species. The final dataset is composed of 14,028 nucleotide sites (4,676 codons).

The time-measured evolutionary history of MeV, RPV, and PPRV was reconstructed using BEAST v1.10.4 (*42*) and the BEAGLE 3 v3.1.2 library to improve computational performance (*45*). Four different evolutionary models, representing an increasing level of complexity, were fitted to the data. All of them used a Yang96 codon substitution model (*46*) and a constant population size tree prior. The first model assumed a strict clock. The second model considered evolution under a strict clock while allowing time-dependency of the nonsynonymous to synonymous substitution rate ratio (dN/dS or ω) under an epoch structure to take long-term purifying selection into account during the phylogenetic reconstruction (*11*). The third model considered evolution under a time-dependent ω, as before, and a fixed local clock (*47*), allowing rate of evolution to change between the PPRV and MeV-RPV clades. The fourth, most complex model considered evolution under a time-dependent ω and a mixed effects clock, where in addition to a fixed local clock, random effects could influence the evolutionary rate on each branch (*48*). The inclusion of the stem of the PPRV clade for the fixed local clock in the third and fourth models were parameterized, allowing the inclusion or not of the stem branch in the model during the inference.

For each model, multiple independent MCMC chains were run until convergence and proper mixing of the model parameters were achieved, as measured by sufficiently high effective sample sizes (ESS > 200) using Tracer v1.7 (*43*). Trees and log files were combined after removing burn-in and posterior distributions of trees were summarized as maximum clade credibility (MCC) trees using TreeAnnotator 1.10.4 (*42*). The time to most recent common ancestor (TMRCA) of the whole tree and the MeV-RPV clade were summarized using mean estimates and 95% highest posterior density (HPD) intervals.

**Supplementary Text**

S1. Directionality of the host switch: from cattle to humans or vice versa?

Cattle domestication, which started about 10,000 years ago (*49*), increased contact between humans and bovines and provided ample opportunity for spill-over of infectious diseases in either direction (*50*). In the case of measles, the predominant view is that the common ancestor of MeV and RPV was a cattle-infecting virus that eventually was transmitted to humans (*6*). MeV and RPV both require large populations to sustain endemicity (i.e. greater than the critical community size (CCS) of 250,000 – 500,000 individuals for measles (*23-25*) and ca. 200,000 for rinderpest (*51*)). The assumption that herds of cattle and wild artiodactyls were sufficiently large to support an RPV-like pathogen long before human populations met the MeV CCS is a central argument to support the hypothesis that measles originated from a cattle-infecting ancestor (*52*, *53*). Indeed, it seems plausible that growing herds of domesticated cattle in combination with wild aurochs and other susceptible artiodactyls had the numbers to support circulation of a multi-host virus like RPV. However, we lack reliable estimates of population sizes of cattle and wild ungulates throughout ancient history which would be required to validate this theory.

Historic sources that may refer to rinderpest exist much earlier than for measles. Descriptions consistent with rinderpest date back as far as the 2$^{nd}$ millenium BCE to an Egyptian veterinary papyrus and can be found on an Indian palm leave text from ca. 1000 BCE as well as in several antique texts (*27*, *54*, *55*). However, as in the case of measles (and in fact most diseases) an unambiguous identification of rinderpest from such historical documents is impossible.

Perhaps the most compelling and supported argument in favor of cattle to human spill-over derives from the phylogeny of the morbillivirus genus. The closest relative to MeV/RPV is PPRV, a virus that mainly infects sheep and goats, but is also known to infect other artiodactyls

(*56*). Phylogenetic proximity facilitates host switches of pathogens (*57*), making a transmission from small ruminants to cattle and then to humans more likely than a switch from small ungulates to humans and then back to cattle (a host switch from humans via small ruminants to cattle can be ruled out based on the new divergence dates). Modern day PPRV causes occasional subclinical infections in cattle (*58*, *59*), showing that barriers to spill-over between small ruminants and cattle are low. It is also conceivable that the common ancestor of PPRV and MeV/RPV infected a broad range of ungulates including cattle and diverged into two more specialized pathogens.

At a deeper evolutionary scale, small mammals appear to be the ancestral hosts of paramyxoviruses (rodents in the case of morbilliviruses; *60*, *61*). Most human-infecting paramyxoviruses appear to have a relatively recent zoonotic origin, including a number of instances where it is known that domesticates have acted as intermediate hosts (e.g. pigs and horses for Nipah and Hendra viruses; *62*, *63*). Thus far, the opposite directionality (human to domesticate) has never been documented. This imbalance might either reflect insufficient discovery effort in domesticates or represent a complex biological trait shared by all paramyxoviruses – the latter possibility rather supporting a cattle to human transmission of the ancestor of MeV and RPV.

All these considerations remain speculative and a scenario of human to cattle transmission, though there is little support for this, cannot be excluded. However, irrespective of the direction of the host switch, the MeV/RPV divergence had enormous consequences on human health, either directly as a major childhood disease or indirectly via its devastating effects on cattle and thereby on human nutrition.

### S2. Pathological report

The pathological report pertaining to the 1912 museum specimen of a 2 year-old child states that the patient was hospitalized with a diagnosis of measles, bronchitis, bronchiolitis and bronchopneumonia. The child died after 3 days in the hospital and a post mortem examination was performed one day after death. The gross pathological findings were translated from German and are listed below.

> "Pathological Diagnosis: Measles-related bronchopneumonia of both lungs with multiple foci of inflammation and mild interstitial pneumonia, bronchitis, tracheitis, bronchiectasis of the left lung, pharyngitis, sub-pleural hemorrhage, alveolar emphysema, swelling of mesenterial and isolated ileal lymph nodes.
>
> The body of the child was well muscled, had a strong built and was well nourished. The skin of the upper chest showed red spots that did not vanish upon pressure. The peritoneum was smooth and glossy. The liver protruded two finger's breadth under the lower end of the costal arch. Position of the diaphragm on both sides: 5$^{th}$ rib. The cartilage-bone junctions of the ribs are distended. After removal of the sternum, the lungs hardly retract into their cavities. The pleural cavities are free of foreign content. The pericardium is filled with a few ccm of clear, serous liquid. Upon opening the heart, uncoagulated blood flows from the heart ventricles and atriums.
>
> The heart itself, about 1.5 times the size of the subject‘s fist, is taut, myocardium left: 1.1 – 0.2 cm, myocardium right: 0.2 cm, of brownish-red color, the endocardium is smooth and glossy, the cardiac valves are thin and delicate. The interior of the heart is filled with blood and postmortem clots.
>
> Left lung: pleura is smooth and glossy, the color varies between a light greyish red and a dark blueish red. The light areas protrude over the surrounding tissue, smaller air bubbles are visible and big sub-pleural bubbles present like a string of pearls. The dark areas feel

dense and rubbery; the light areas are air-filled and elastic. There are several red spots under the pleura that do not vanish upon pressure. The cut surface is very uneven; the color is light mottled with a dark reddish blue. The liquid that escapes the tissue contains only very little air and is bloody. In case of the smallest bronchi, there is a small amount of yellowish green liquid. Some bronchi with more pronounced reddening of the mucosa are also filled with greenish yellow mucus. Isolated bronchi show cylindrical dilations. Pulmonary lymph nodes are swollen to the size of hazelnuts, are very firm and extremely red.

Right lung: findings correspond to left lung.

Throat: the pharyngeal mucosa and the area around the larynx opening are reddened. Both tonsils are slightly swollen and filled with a small amount of greenish yellow liquid that spills out when tonsils are cut. Esophagus no particular findings. The tracheal mucosa is extremely red, the lymph nodes of the neck are swollen and very red. Thyroid gland, thoracic aorta without particular findings.

Spleen: 7.5 x 4.5 x 2.5 cm. The color is blue-red, the consistency firm-elastic. The surface is smooth and glossy. The cut surface clearly shows the structure of the lymph follicles, the color is blue-red, the pulpa is intact.

Kidneys: (2.5 x 3 x 3 cm) capsule can be removed easily, color grey-red, surface smooth and glossy. Consistency: firm-elastic. On the cut surface cortex and medulla can easily be distinguished. The parenchyma is slightly cloudy, renal pelvis no particular findings. Right kidney (7.5 x 3 x 2.5 cm): findings correspond to left kidney. Adrenal glands are very pale, otherwise without particular findings.

Urinary bladder, genital tract, rectum without particular findings.

Liver: (16.5 x 11 x 5.5 cm) surface is smooth and glossy. Color: brown-red. Consistency: firm-elastic. On the cut surface the hepatic lobules are completely clear, the parenchyma is clear, blood content is moderate. Gall bladder no particular findings.

Stomach, pancreas, duodenum without particular findings. Some ileal lymph nodes are slightly swollen. Abdominal aorta no particular findings.”

S3. Histopathological report

We performed histopathological analyses of the 1912 formalin-fixed lung specimen. For conventional histopathology, slides of lung tissue were stained with hematoxylin and eosin (fig. S1). We observed an active, focally purulent bronchopneumonia with widespread interstitial pneumonia, associated with multinucleated giant cells of the Warthin-Finkeldey cell type. These findings, while not specific for MeV infection, are consistent with measles-related bronchopneumonia. Immunohistochemical approaches were unsuccessful due to the age and low quality of the specimen.

S4. Phylogenetic exploration of the 1912 MeV sample genome sequence

The Bayesian phylogenetic tree placed the 1912 genome basal to all modern genomes; the two genomes from 1960 clustered together with the Edmonston strain isolated in 1954 (fig. S2). To confirm the phylogenetic placement of the 1912 genome, we explored how this sample branched in a full-genome MeV maximum likelihood phylogeny without date information. On an unrooted phylogeny, the 1912 sample branched very deeply in the tree, on the branch linking the H1 genotype to the others (fig. S3A). A similar tree, without the 1912 sample and optimally rooted to minimize residual mean squared distance of the sampling time and root-to-tip divergence correlation in TempEst, shows that the same branch is basal to the whole phylogeny (fig. S3B), confirming the phylogenetic placement of the 1912 sample as basal to all more recent samples. This is further supported by the strong correlation, computed without the 1912 sample, between sampling time and root-to-tip divergence (tree rooted with the 1912 sample, rooting optimized to minimize residual mean squared distance) for the complete genome dataset of MeV (n= 130), including the 1912 sample ($R^2 = 0.624$, $p < 2.2 \times 10^{-16}$; fig. S3C).

S5. Model comparison for divergence dating of MeV and RPV

The four models tested, representing an increasing level of complexity, yielded different results in their model parameter estimates (table S2, Fig. 2). The general trend is that the more complex the model, the more ancient the divergence time estimate is and the wider the 95% HPD interval, for both MeV and RPV and PPRV and MeV/RPV divergence time estimates. The models are essentially a range of nested models, for which new parameters in a more complex model accommodate an additional source of rate variation relative to the simpler model. In each case, the impact on the parameter estimates implies that the contribution of the additional source of rate variation is significant. The most complex model, combining a codon substitution model with time-dependent $\omega$ and a mixed-effect clock model (associating fixed local clock for MeV/RPV and PPRV, and random effects on the evolutionary rate) therefore provides the best fit to the data (table S2). The branch ancestral to the PPRV clade did not receive any support for sharing the same fixed effect modelled within the PPRV clade.

To investigate the impact of the new 1912 MeV genome's date, an additional model, based on the most complex model we previously assessed (codon substitution model with time-dependent $\omega$ and a mixed-effect clock model) was set up. In this model, the 1912 MeV genome's date was considered as unknown and estimated from the data. This model did not yield substantially different estimates (mean: 300 BCE; 95% HPD interval [-1021; 345]), suggesting the 1912 genome's sampling date had no particular impact on the reconstruction of the deep evolutionary history of the three viruses.

S6. Estimating historical urban populations

We have obtained estimates of the past population of cities from standard historical and archaeological literature, collating data from Inoue et al. 2015 (*26*) and Morris 2013 (*29*), who in turn draw from Chandler 1987 (*64*), Modelski 2003 (*65*), Bairoch 1988 (*66*), and other more localized studies. Estimating historical urban populations entails considerable methodological and empirical difficulties. The very definition of a settlement is a complex question. Inoue et al. (*26*) define a settlement as a "spatially contiguous built-up area," and this definition reasonably allows us to capture what is most epidemiologically important, the number of people living in contact within a single area. Ultimately, estimates for the scale of premodern urban populations are based on one of two kinds of data: written or archaeological. Written evidence may include literary observations, travelers' reports, censuses, or references to the amount of food distributed in a city; obviously the reliability of such evidence varies. Archaeological evidence can be based on the size of city walls or the extent of the built-up area of a settlement, as determined by excavation or field survey. Even if the spatial extent of a settlement is known, an estimate of the population density is required to extrapolate the total population. Settlement densities varied across time and space, and estimates are highly sensitive to assumptions about the nature of an urban settlement. Most often, settlement densities are estimated at 100-200 persons per hectare, even though lower and much higher densities are attested. An overview of the methodological issues is provided by Pasciuti and Chase-Dunn 2002 (*67*).

Despite the imprecision and uncertainty involved in estimating premodern urban populations, we note the extreme consensus on the issue of importance to this study. No credible scholarship maintains that any urban population passed the CCS for the MeV prior to 500 BCE. Simply, cities of the Neolithic, Bronze Age, and early Iron Age could not acquire the size necessary to sustain endemic measles infection. Thereafter, however, from the late first millennium BCE, technologies (both economic and political) crossed a threshold allowing larger populations, and from this period onward, there were consistently human populations above the CCS for MeV.

We also observe that the concept of a "critical community size" was developed to model the epidemiological dynamics of an infectious disease in a closed or isolated population such as an island. MeV, because it is a human specialist that causes an acute infection conferring strong and lasting immunity on survivors, has always been a paradigm for the study of critical community size. Yet CCS provides a way to predict local, rather than global, extinction; connectivity between settlements could create a larger effective number of susceptible individuals, buffering against extinction. Nevertheless, given the speed with which measles epidemics occur, and the efficacy of the acquired immunity, epidemiologists have held that urban populations must have passed the CCS or close to it for MeV, in its human-specialized form, to have evolved and avoided extinction (*68*).

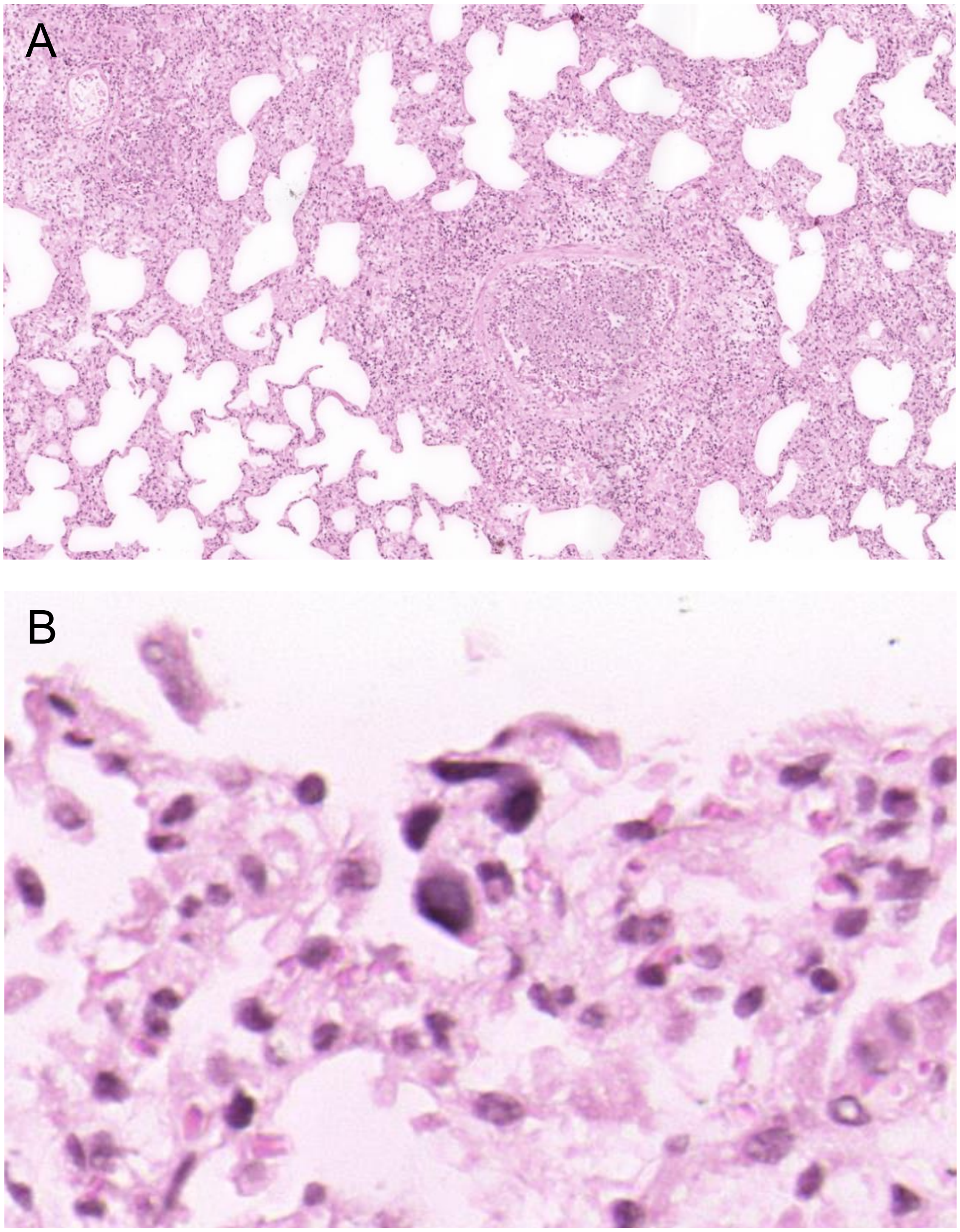


**Fig. S1. Histopathology of formalin fixed lung.** Hematoxylin and eosin stain. Magnified 50x (A) and 400x (B). Active, focally purulent bronchopneumonia with widespread interstitial pneumonia, associated with multinucleated giant cells of the Warthin-Finkeldey cell type.

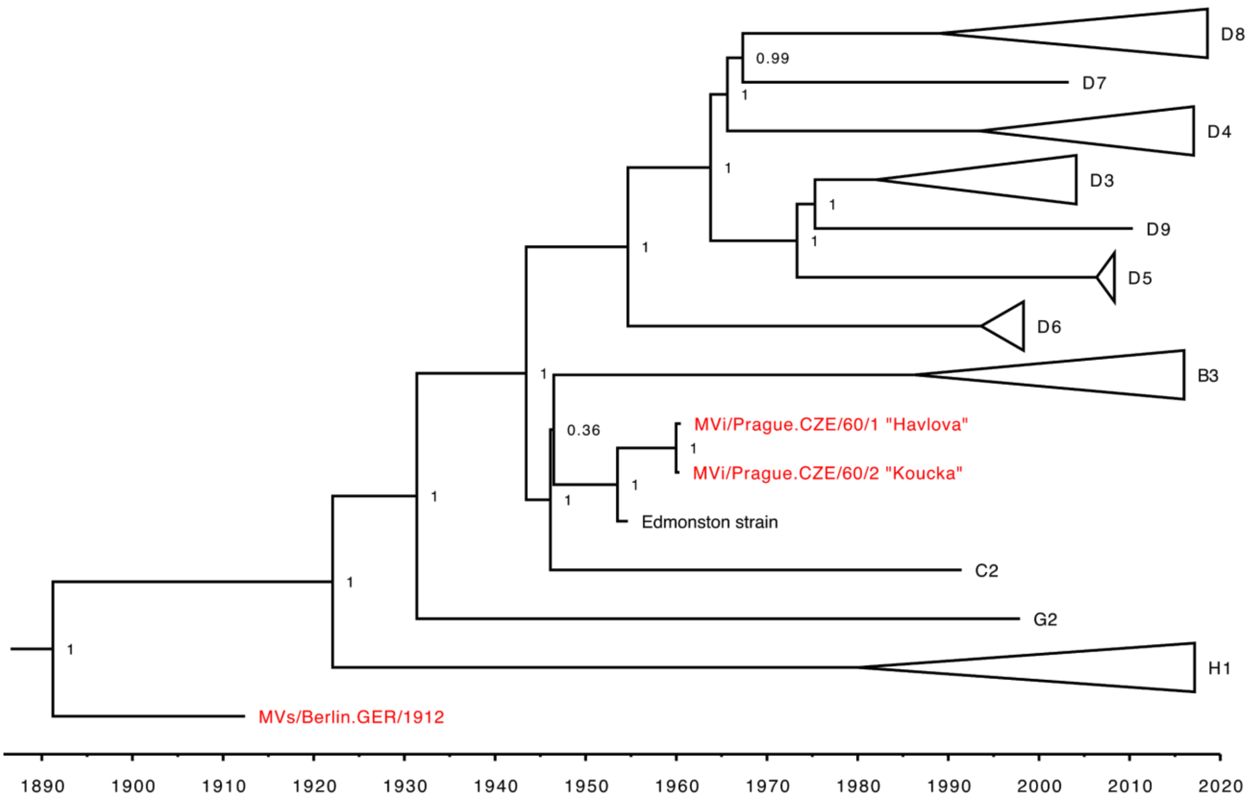


**Fig. S2. Timed phylogenetic tree of Measles virus (n=130) including the three sequences generated in this study (highlighted in red), showing the main genotypes.** This tree is the maximum clade credibility (MCC) tree derived from a Bayesian evolutionary inference for 130 full genomes of MeV (including the 3 new ones highlighted in red). A time scale is shown at the bottom. Numbers represent node posterior probability support.

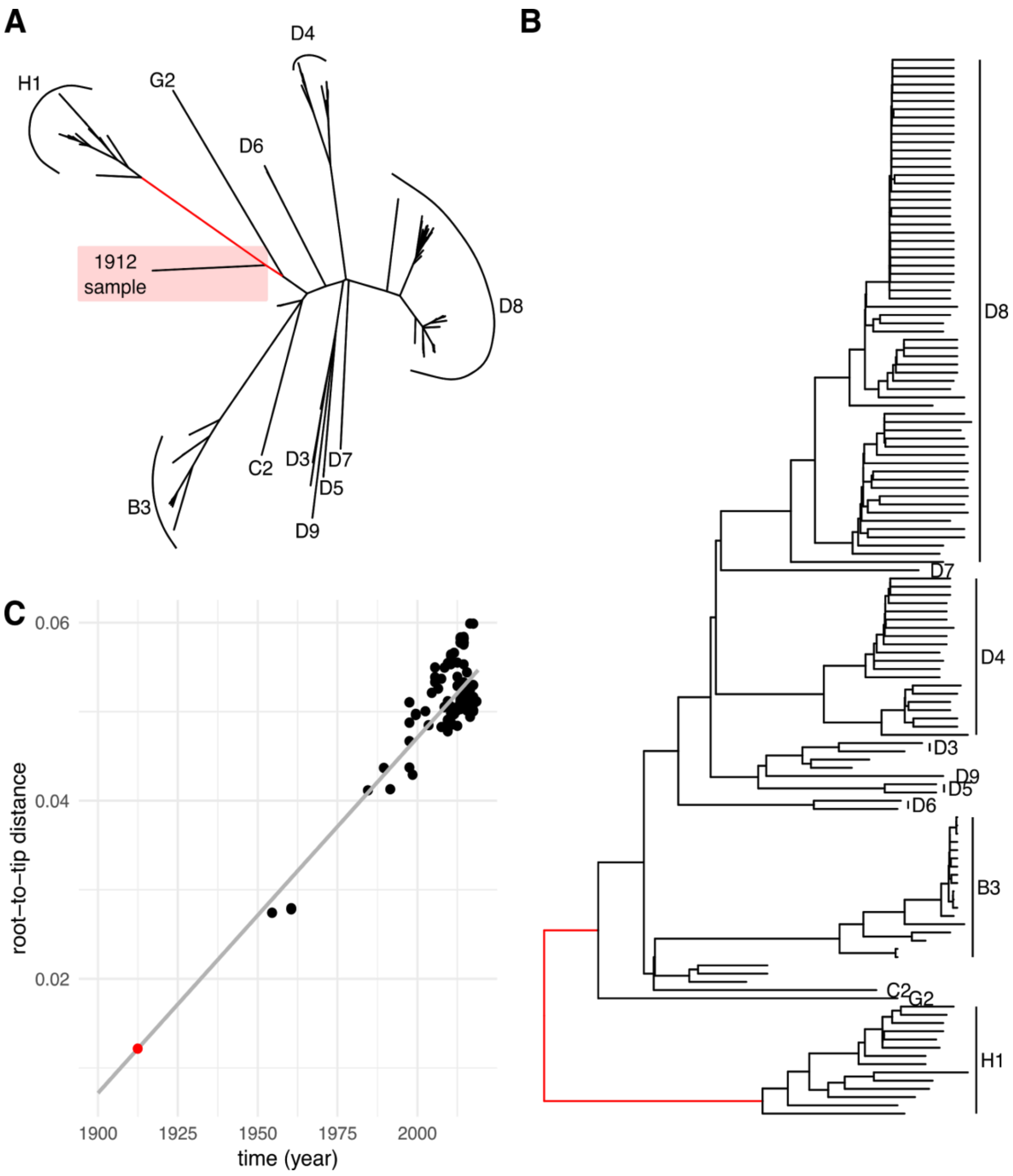


**Fig. S3. Phylogenetic exploration of the 1912 measles genome sequence from Germany.** (A) Unrooted tree for all 130 MeV including the 1912 measles genome (red background). The branch the 1912 roots in is highlighted in red (B) Rooted time-tree for 129 MeV (excluding 1912) genomes. Root point is optimized to minimize residual mean squared distance of the sampling time and root-to-tip divergence correlation in TempEst. The branch where the 1912 genome roots in the unrooted tree is highlighted in red. (C) Correlation between sampling time and root-to-tip divergence for the complete genome dataset of MeV (n= 130). Rooting is optimized to minimize residual mean squared distance of the sampling time and root-to-tip divergence correlation in TempEst. The correlation line (in gray) has been computed without the 1912 genome. The 1912 genome is highlighted in red.

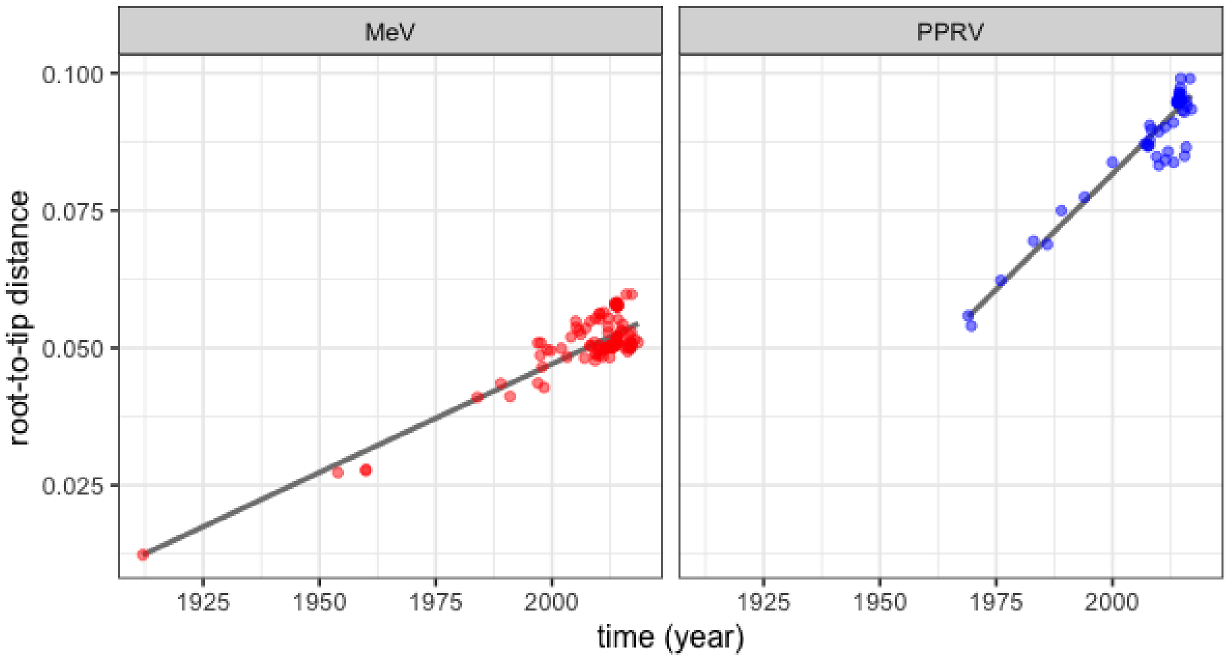


**Fig. S4. Correlation between sampling time and root-to-tip divergence for the complete genome dataset of MeV (n= 130) in red and PPRV (n=65) in blue**. Rooting is optimized to minimize residual mean squared distance of the sampling time and root-to-tip divergence correlation in TempEst.

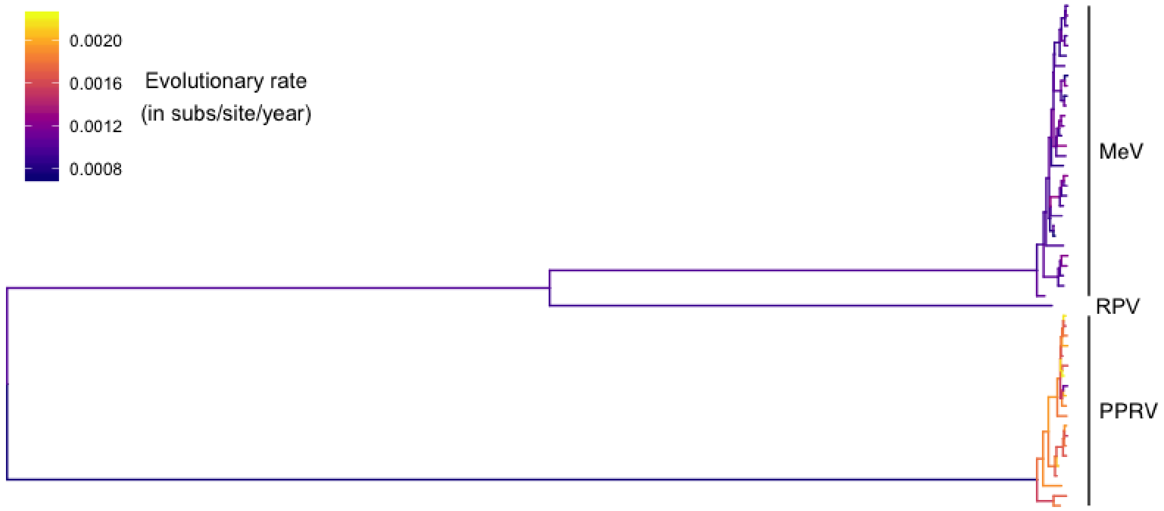


**Fig. S5. Evolutionary rates (in substitutions/site/year) across branches under a mixed effects clock and time-dependent ω.**

| GenBank accession number | Date (in decimal year) | Uncertainty (prior) | Virus |
|---|---|---|---|
| XXX (655_3reads_50_1912) | 1912.426 | - | MeV |
| XXX (6114_3reads_50_1960) | 1960 | 1 year (uniform) | MeV |
| AF266288 | 1954 | 1 year (uniform) | MeV |
| AB481087 | 1989 | 1 year (uniform) | MeV |
| MG912589 | 1991 | 1 year (uniform) | MeV |
| DQ227319 | 1997 | 1 year (uniform) | MeV |
| MG912590 | 1997.62739726027 | - | MeV |
| MG912594 | 1997.49315068493 | - | MeV |
| GQ376026 | 1999.69863013699 | - | MeV |
| KJ755974 | 1999 | 1 year (uniform) | MeV |
| JN635410 | 2003.28219178082 | - | MeV |
| JN635409 | 2004.12568306011 | - | MeV |
| JN635406 | 2008.18852459016 | - | MeV |
| MH356248 | 2009.10684931507 | - | MeV |
| KY969476 | 2010.35342465753 | - | MeV |
| KY969479 | 2013.87945205479 | - | MeV |
| MG912591 | 1997.91506849315 | - | MeV |
| MH356249 | 2014.20547945205 | - | MeV |
| MH356237 | 2016.69672131148 | - | MeV |
| MH356243 | 2016.77322404372 | - | MeV |
| MH356238 | 2017.07945205479 | - | MeV |
| MH356240 | 2017.02191780822 | - | MeV |

| | | | |
|---|---|---|---|
| KT588921 | 2013.53424657534 | - | MeV |
| KC117298 | 2010.21917808219 | - | MeV |
| MG972194 | 2017.17534246575 | - | MeV |
| KT732214 | 2012.1174863388 | - | MeV |
| KT732224 | 2014.14794520548 | - | MeV |
| KX838946 | 2016.00819672131 | - | MeV |
| KT732261 | 2013.55342465753 | - | MeV |
| MH173047 | 2017.84657534247 | - | MeV |
| X98291 | 1910 | (exponential prior, mean 10 year, offset 57.84658 years)* | RPV |
| KR781450 | 1969 | 1 year (uniform) | PPRV |
| EU267274 | 1976 | 1 year (uniform) | PPRV |
| KJ466104 | 2010 | 1 year (uniform) | PPRV |
| KU236379 | 2015.51506849315 | - | PPRV |
| MF741712 | 2011.95890410959 | - | PPRV |
| KR781449 | 2011.38356164384 | - | PPRV |
| KR140086 | 1994 | 1 year (uniform) | PPRV |
| JX217850 | 2008 | 1 year (uniform) | PPRV |
| MG581412 | 2008.3306010929 | - | PPRV |
| KR261605 | 2014.70684931507 | - | PPRV |
| KY888168 | 2016.66666666667 | - | PPRV |
| NC_006383 | 2000 | 1 year (uniform) | PPRV |
| KC594074 | 2008 | 1 year (uniform) | PPRV |
| KY885100 | 2015.8602739726 | - | PPRV |
| KJ867541 | 2010 | 1 year (uniform) | PPRV |

| | | | |
|---|---|---|---|
| MF678816 | 2017 | 1 year (uniform) | PPRV |
| KR828813 | 2013.12328767123 | - | PPRV |
| EU267273 | 1989 | 1 year (uniform) | PPRV |
| KJ867544 | 1983 | 1 year (uniform) | PPRV |
| KM463083 | 2011.32876712329 | - | PPRV |

**Table S1. Accession numbers of sequences used in the Bayesian phylogenetic analyses performed to date MeV/RPV/PPRV divergence.** * This sequence derives from a field isolate from 1910 that was then passaged until the late 1950's (*69*). To model its dating uncertainty, we used an exponential prior with an offset of 57.84658 years (date of the most recent sequence minus 1960) and a mean of 10.

| **Model** | **Parameters** | **Mean** | **95% HPD** |
|---|---|---|---|
| Strict clock | Ω | 0.084 | [0.081; 0.088] |
| | Clock rate *r* | $1.275 \times 10^{-3}$ | [$1.207 \times 10^{-3}$; $1.347 \times 10^{-3}$] |
| | Divergence $date_{MeV - RPV}$ | 938 | [862; 1014] |
| | Divergence $date_{PPRV - MeV/RPV}$ | 451 | [345; 559] |
| Strict clock and time-dependent ω | ω: Intercept $\beta 0$ | -1.296 | [-1.440, -1.146] |
| | ω: Slope $\beta 1$ | -0.284 | [-0.321, -0.248] |
| | Clock rate *r* | $1.268 \times 10^{-3}$ | [$1.188 \times 10^{-3}$; $1.345 \times 10^{-3}$] |
| | Divergence $date_{MeV - RPV}$ | 315 | [64; 523] |
| | Divergence $date_{PPRV - MeV/RPV}$ | -668 | [-1178; -307] |
| Fixed local clock and time-dependent ω | ω: Intercept $\beta 0$ | -1.217 | [-1.345, -1.096] |
| | ω: Slope $\beta 1$ | -0.309 | [-0.338, -0.280] |
| | Clock rate $r_{MeV/RPV}$ | $1.111 \times 10^{-3}$ | [$1.023 \times 10^{-3}$; $1.191 \times 10^{-3}$] |
| | Clock rate $r_{PPRV}$ | $1.688 \times 10^{-3}$ | [$1.558 \times 10^{-3}$; $1.823 \times 10^{-3}$] |
| | Divergence $date_{MeV - RPV}$ | -104 | [-349; 112] |
| | Divergence $date_{PPRV - MeV/RPV}$ | -1570 | [-2191; -1024] |
| Mixed-effect clock and time-dependent ω | ω: Intercept $\beta 0$ | -1.259 | [-1.389; -1.132] |
| | ω: Slope $\beta 1$ | -0.292 | [-0.322; -0.260] |

| | | | |
|---|---|---|---|
| | Clock rate $r_{MeV/RPV}$ | $0.980\times 10^{-3}$ | [$0.824 \times 10^{-3}$; $1.160 \times 10^{-3}$] |
| | Clock rate $r_{PPRV}$ | $1.727 \times 10^{-3}$ | [$1.419 \times 10^{-3}$; $2.107 \times 10^{-3}$] |
| | Clock rate *sd* | 0.208 | [0.144; 0.276] |
| | Divergence $date_{MeV - RPV}$ | -345 | [-1066; 319] |
| | Divergence $date_{PPRV - MeV/RPV}$ | -2821 | [-4177; -1665] |
| Mixed-effect clock and time-dependent ω, with 1912 date sampled | ω: Intercept *β0* | -1.261 | [-1.389, -1.113] |
| | ω: Slope *β1* | -0.292 | [-0.323, -0.261] |
| | Clock rate $r_{MeV/RPV}$ | $0.990 \times 10^{-3}$ | [$0.808 \times 10^{-3}$; $1.189 \times 10^{-3}$] |
| | Clock rate $r_{PPRV}$ | $1.731 \times 10^{-3}$ | [$1.438 \times 10^{-3}$; $2.083 \times 10^{-3}$] |
| | Clock rate *sd* | 0.207 | [0.145, 0.275] |
| | Divergence $date_{MeV - RPV}$ | -300 | [-1021; 345] |
| | Divergence $date_{PPRV - MeV/RPV}$ | -2813 | [-4185; -1599] |
| | $Date_{1912\ sample}$ | 1923 | [1883; 1962] |

**Table S2. Parameters estimates for each model tested.**

34. G. Jäger, ClipAndMerge. *https://github.com/apeltzer/ClipAndMerge*.

35. B. Institute, Picard. *http://broadinstitute.github.io/picard*.

36. A. Peltzer, DeDup. *https://github.com/apeltzer/DeDup*.

37. Geneious 11.1.5. *https://www.geneious.com*.

45. D. L. Ayres, M. P. Cummings, G. Baele, A. E. Darling, P. O. Lewis, D. L. Swofford, J. P. Huelsenbeck, P. Lemey, A. Rambaut, M. A. Suchard, BEAGLE 3: Improved Performance, Scaling, and Usability for a High-Performance Computing Library for Statistical Phylogenetics. *Systematic biology,* (2019).

53. W.H. McNeill, Plagues and Peoples (Anchor Press, 1976)

54. P.-P. Patoret, K. Yamanouchi, U. Mueller-Doblies, M. M. Rweyemamu, M. Horzinek, T. Barrett, "Rinderpest – an old and worldwide story: history to c.1902" in *Rinderpest and Peste des Petits Ruminants: Virus Plagues of Large and Small Ruminants*, T. Barrett, P.-P. Pastoret, W. Taylor, Eds. (Elsevier, 2005).

55. J. Blancou, "Old prophylactic methods" in *Rinderpest and Peste des Petits Ruminants: Virus Plagues of Large and Small Ruminants*, T. Barrett, P.-P. Pastoret, W. Taylor, Eds. (Elsevier, 2005).

56. OIE, Peste des Petits Ruminants https://www.oie.int/fileadmin/Home/eng/Animal_Health_in_the_World/docs/pdf/Disease_cards/PESTE_DES_PETITS_RUMINANTS.pdf

64. T. Chandler, *Four thousand years of urban growth: an historical census* (Lewiston, 1987).

65. G. Modelski, *World Cities: -3000 to 2000* (Washington, D.C., 2003).

66. P. Bairoch, *Cities and economic development from the dawn of history to the present* (Chicago, 1988).

67. D. Pasciuti, C. Chase-Dunn, "Estimating The Population Sizes of Cities," 2002. https://irows.ucr.edu/research/citemp/estcit/estcit.htm Accessed 10/25/2019.

68. A. Cliff, P. Haggett, and M. Smallman-Raynor, *Measles: an historical geography of a major human viral disease: from global expansion to local retreat, 1840-1990* (Oxford, 1993).

69. M. D. Baron, T. Barrett, Rescue of rinderpest virus from cloned cDNA. Journal of virology 71, 1265-1271 (1997).
